# A single extracellular vesicle-based platform supporting both RBD protein and mRNA vaccination against SARS-CoV-2

**DOI:** 10.1101/2025.06.23.661193

**Authors:** Jihwa Chung, Kyoung Hwa Kim, Shung Hyun An, DaeHo Bae, Jae Hwan Kim, Kihwan Kwon, Seok-Hyun Kim

**Affiliations:** Exollence Co., Ltd., Seoul 07985, Republic of Korea; Department of Internal Medicine, Cardiology Division, School of Medicine, Ewha Womans University, Seoul 07985, Republic of Korea

**Keywords:** Extracellular vesicles, SARS-CoV-2, RBD antigen, mRNA vaccine, protein subunit vaccine, non-LNP delivery, shock wave–mediated post-loading, SWEET platform

## Abstract

The coronavirus disease 2019 (COVID-19) pandemic caused by severe acute respiratory syndrome coronavirus 2 (SARS-CoV-2) has highlighted the need for novel, effective, and safe vaccine platforms. Extracellular vesicles (EVs) facilitate cell-to-cell communication and can modulate immune responses by delivering antigens. Here, we propose a versatile EV-based vaccine plat-form in which immune-stimulating EVs, engineered using acoustic shock waves (SW), are used to deliver either protein or mRNA antigens. Immunostimulatory EVs derived from LPS-activated THP-1 monocytes (aTEVs) were loaded with SARS-CoV-2 receptor-binding domain (RBD) protein or RBD-encoding mRNA via SW post-loading. Both aTEV-RBD-Pro and aTEV-RBD-mRNA elicited robust RBD-specific humoral and cellular immune responses in mice without the need for external adjuvants. Moreover, lyophilized aTEV-RBD-Pro vaccines retained immunogenicity after storage, supporting their chain-independent stability. These results demonstrate that a single EV-based platform can independently support both protein- and mRNA-based vaccination, highlighting its flexibility for next-generation vaccine development.

## Introduction

The coronavirus disease 2019 (COVID-19) pandemic caused by severe acute respiratory syndrome coronavirus 2 (SARS-CoV-2) has affected hundreds of millions of people worldwide and caused millions of deaths. This crisis has highlighted the urgent need for novel vaccine platforms that are safe, effective, and rapidly adaptable to emerging pathogens. Among various approaches, mRNA-based vaccines have demonstrated remarkable efficacy and scalability [1, 2], while nanocarrier-based systems such as lipid nanoparticles and microparticles have also shown clinical promise [3]. However, current nanocarrier platforms still face limitations, including potential immunogenicity, variable adjuvant responses, and concerns over long-term safety and durability [4, 5].

As an alternative, extracellular vesicles (EVs) have emerged as natural, biocompatible carriers with intrinsic immunomodulatory properties [6, 7]. EVs are secreted by nearly all cell types, range in size from 20 to 4000 nm, and carry a diverse cargo of proteins, lipids, and nucleic acids [8]. Their inherent ability to interface with immune cells makes them attractive candidates for vaccine delivery. Notably, EVs derived from mesenchymal stem cells (MSCs) or immune cells such as M1 macrophages and activated THP-1 cells have demonstrated immunostimulatory functions in various preclinical models, including those involving SARS-CoV-2 and cancer [6, 9, 10]. Exosome-based vaccines have been studied since the early 2000s and shown to elicit antigen-specific cytotoxic T-cell responses [11–13].

Although EVs are promising delivery vehicles, efficient encapsulation of functional cargo— particularly large proteins or mRNAs—remains a technical challenge. Conventional EV loading strategies such as incubation, transfection, electroporation, and extrusion often result in low efficiency or vesicle damage [14–18]. Therefore, there is a pressing need for safer and more robust EV engineering methods.

Shock wave (SW) technology, a high-energy acoustic pulse previously used in lithotripsy and musculoskeletal therapy [19, 20], has recently been explored as a means to enhance membrane permeability and facilitate molecular delivery into biological systems [21–23]. In our earlier work, we demonstrated that low-energy SW could enable direct transfection of siRNAs into mammalian cells via a novel physical mechanism [24]. Building on this principle, we later developed a scalable and efficient EV engineering method termed SWEET (Shock Wave Extracellular vesicle Engineering Technology), which enables high-efficiency encapsulation of diverse biological cargos—including nucleic acids such as siRNA and mRNA, as well as proteins—into EVs without com-promising vesicle integrity. In a recent study, we reported that SWEET-based siRNA-loaded EVs effectively suppressed KRAS-mutant tumor growth in vivo, highlighting the platform’s translational potential [25].

Building on this platform, we applied SWEET to encapsulate SARS-CoV-2 receptor-binding domain (RBD) protein and RBD-encoding mRNA into immunostimulatory EVs derived from LPS-activated THP-1 cells (aTEVs). We show that both aTEV-RBD-Pro and aTEV-RBD-mRNA elicited robust RBD-specific humoral and cellular immune responses in mice, without the need for additional adjuvants. Furthermore, lyophilized aTEV vaccines retained their immunogenicity, supporting the possibility of cold chain–free distribution. These findings demonstrate that a single engineered EV platform can independently support both protein and mRNA vaccination, offering a flexible and scalable approach to future vaccine development.

## Methods

### Cell culture

THP-1 human monocytes were obtained from the Korean Cell Line Bank (Seoul, South Korea) and were cultured in RPMI 1640 (Welgene) supplemented with 10% fetal bovine serum (FBS; Biowest) and 1% penicillin/streptomycin (Corning). JAWSII Mouse dendritic cells were obtained from the ATCC and were maintained in Alpha Minimal Essential Medium (MEM-α) (Welgene), 20% FBS, and 5 ng/mL recombinant mouse granulocyte-macrophage colony-stimulating factor (GM-CSF; BioLegend), and 1% peni-cillin/streptomycin (Corning). The cells were cultured in a 5% CO2 atmosphere at 37°C

### Isolation and purification of naïve TEVs (nTEVs) and aTEVs

To produce aTEVs, THP-1 cells were stimulated with lipopolysaccharide (LPS; from Escherichia coli strain O111:B4, Sigma), following a protocol adapted from Jesus et al. [10]. Briefly, cell culture medium was supplemented with 1% EV-free FBS, which had been centrifuged at 100,000 g for 16 h at 4°C, to eliminate contaminating bovine serum-derived EVs. THP-1 cells were incubated with or without 100 ng/mL LPS for 16 h at a density of 8 × 10^5^/mL. The culture medium was centrifuged at 300 × g for 5 min at RT to remove cell debris. The supernatant was passed through a 0.22 μm filter and concentrated by tangential flow filtration (Vivaflow 50, 100 kDa MWCO; Sartorius). nTEVs and aTEVs were resuspended in 50 mM trehalose-PBS and stored at –80 C. To prepare nTEV and aTEV supernatants, each was centrifuged at 100,000 × g for 2 h at 4°C.

### Endotoxin assay (LAL test)

Residual endotoxin levels in the final aTEV preparations were assessed using a Limulus Amebocyte Lysate (LAL) chromogenic assay kit (ToxinSensor™, GenScript), following the manufacturer’s instructions. Absorbance was measured at 405 nm and endotoxin concentrations were calculated using a standard curve. LAL assay was per-formed on the resuspended EV pellet after final purification.

### Nanosight tracking analysis

The size and concentration of purified nTEVs or aTEVs were analyzed using the ZetaView Nanoparticle Tracking Analyzer (Particle Metrix, Germany) with a 488 nm laser light source. Prior to measurements, the accuracy of the Zetaview instrument was confirmed by conducting measurements using a 100 nm standard nanosphere. Samples were diluted in PBS to 1:1000 to 1:5000 to a particle count of 200 to 500.

### Cryo-electron microscopy (Cryo-EM)

aTEVs, with or without SW treatment, were applied to glow-discharged 300 mesh Quantifoil holey carbon EM grids. The grids were vitrified using a Vitrobot Mark IV (FEI Company) at 4°C and 100% humidity. Excess sample was removed by blotting once with filter paper, and the blotted grids were dipped in liquid ethane. Cryo-EM imaging was performed on a 300 kV Titan Krios electron microscope (FEI Company).

### Western blotting

Cells or aTEVs were lysed in radioimmunoprecipitation assay (RIPA) buffer. The protein concentrations of the lysates were measured using a bicinchoninic acid protein assay (BCA) (Thermo Fisher Scientific) following the manufacturer’s instructions. The lysates were separated by sodium dodecyl sulfate–polyacrylamide gel electrophoresis (SDS-PAGE). Protein bands were transferred to nitrocellulose membranes (Amersham Biosciences). After blocking with 3% bovine serum albumin in Tris-buffered saline with Tween 20 at room temperature (RT), the membranes were incubated with the primary antibodies at 4°C overnight. Next, they were incubated with secondary antibodies for 1 h at RT and visualized by chemiluminescence (Thermo Fisher Scientific). The primary antibodies were anti-CD9 (Cell Signaling Technology), anti-CD63 (Abcam), anti-CD81 (Santa Cruz Biotechnology), anti-syntenin-1 (Genetex), anti-Alix (Cell Signaling Tech-nology), anti-TSG101 (Santa Cruz Biotechnology), anti-flotillin-1 (Abcam), anti-calnexin (Merck), anti-GM130 (Cell Signaling Technology), anti-RBD (R&D Systems), and anti-GAPDH (Santa Cruz Biotechnology).

### RT-PCR Analysis

LPS-stimulated THP-1 cells were evaluated for the expression of marker genes of pro-inflammatory M1-polarized macrophages. Total RNA was extracted using QIAzol Lysis Reagent (Qiagen) according to the manufacturer’s instructions. Extracted RNA was reverse-transcribed into cDNA using the Promega GoScript™ Reverse Transcription System (Promega). PCR was carried out in an Applied Biosystems™ QuantStudio™ 3 Real-Time PCR System (Thermo Fisher Scientific) using THUNDERBIRDTM SYBRTM qPCR Mix (Toyobo, Osaka, Japan). Relative mRNA levels were determined by real-time quantitative PCR and normalized to the internal control, GAPDH. The experiments were performed in triplicate. The sequences of the primers were as follows: Human IL-6: forward 5’- AGGAGACTTGCCTGGTGAAA-3’, reverse 5’- CAGGGGTGGTTATTGCATCT-3’; Human IL-1β: forward 5’- GGGCCTCAAGGAAAAGAATC-3’, reverse 5’- TTCTGCTTGAGAGGTGCTGA-3’; Human IL-8: forward 5’- ACTGAGAGTGATTGAGAGTGGAC-3’, reverse 5’- AACCCTCTGCACCCAGTTTTC-3’; Human TNF-α: forward 5’- GAGGCCAA-GCCCTGGTATG-3’, reverse 5’- CGGGCCGATTGATCTCAGC-3’; Human iNOS: for-ward 5’- AGGGACAAGCCTACCCCTC-3’, reverse 5’- CTCATCTCCCGTCAGTT-GGT-3’; Human GAPDH: forward 5’- GAGTCAACGGATTTGGTCGT-3’, reverse 5’- TTGATTTTGGAGGGATCTCG-3’.

### Cargo loading into EVs using SWEET (Shock Wave Extracellular vesicle Engineering Tech-nology)

Cargo encapsulation was performed using SWEET, an acoustic shockwave-based method optimized for post-loading of biologics into extracellular vesicles, as previously [25].

Encapsulation of RBD protein : Recombinant SARS-CoV-2 RBD protein with the Fc region of mouse IgG2a at the C-terminus (RBD-Fc) was purchased from AbClon (Seoul, South Korea). Purified aTEVs were mixed with recombinant SARS-CoV-2 RBD protein in PBS. The mixture was placed in a degassed acoustic chamber and exposed to shock waves at 0.3 mJ/mm² for 1000 pulses. Following treatment, the sample was filtered through a 100 kDa MWCO ultrafiltration column (Amicon Ultra) to remove unencapsulated protein.

Encapsulation of RBD mRNA : RBD-encoding mRNA (1 or 10 µg; RNagene, Republic of Korea) was incubated with aTEVs under the same SWEET conditions (0.3 mJ/mm², 1000 pulses). Post-SW, the mixture was processed through a 100 kDa MWCO filter. For functional validation, HCT116 cells were treated with mRNA-loaded aTEVs, and RBD protein expression was assessed via Western blotting.

### Verification of encapsulated RBD in aTEVs

Quantification of encapsulated RBD protein : To determine the amount of SARS-CoV-2 RBD protein encapsulated in aTEVs, aTEV-RBD-Pro samples were lysed in PBS containing 0.25% Triton X-100 for 2 h at 4 °C on a shaker. The encapsulated protein content was then quantified using a mouse IgG2a ELISA Kit (Invitrogen) following the manufacturer’s instructions with slight modifications. ELISA plates were coated with capture antibody and blocked with PBS containing 0.1% Tween 20 and 1% BSA for 2 h at room temperature. After washing, standards or lysed EV samples and detection antibody were added and incubated for 3 h. TMB substrate (Invitrogen) was used for color development and stopped with stop solution (R&D Systems). Absorbance at 450 nm was measured using a Synergy H1 Microplate Reader (BioTek), and the RBD concentration was determined from a standard curve.

Verification of encapsulated RBD mRNA : To evaluate mRNA incorporation into EVs, the EV–mRNA mixture was subjected to centrifugal filtration using a 100 kDa molecular weight cut-off filter (Amicon Ultra, Millipore). The resulting filtrate and retentate fractions were analyzed by agarose gel electrophoresis. In a separate experiment, Benzonase nuclease (50 U/mL; Sigma) was added to EV preparations and incubated for 30 minutes at 37 °C to remove non-encapsulated mRNA. RNA was then extracted from EVs using TRIzol reagent (Invitrogen) according to the manufacturer’s instructions. The amount of protected RBD mRNA was subsequently measured by reverse transcription followed by quantitative PCR (RT-qPCR) using specific primers targeting the RBD sequence.

### In vivo imaging and tracking of aTEVs

To label aTEVs with DiR, a working solution of XenoLight DiR (320 µg/mL; PerkinElmer) was first prepared according to the manufacturer’s instructions. The DiR solution was then mixed with the appropriate amount of aTEVs and incubated at room temperature for 30 minutes in the dark. Following incubation, unbound dye was re-moved using a 100 kDa Amicon Ultra centrifugal filter unit (Millipore) by centrifugation at 3,000 × g for 10–15 minutes. The resulting DiR-labeled aTEVs were collected and used for subsequent experiments. For in vivo imaging, DiR-labeled aTEVs (50 μg) were injected subcutaneously into the right flank of mice, and their distribution was visualized ex vivo using VISQUE® InVivo Smart-LF (Vieworks, Anyang, South Korea). Inguinal and axillary lymph nodes from the injection and contralateral sides were collected at 24 h post-injection. To identify the cells that took up aTEVs, EVs (50 μg) labeled with red fluorescence (PKH26) were injected subcutaneously. Lymph nodes were collected 24 h after injection and treated with collagenase (2 mg/mL) and DNaseI (40 U/mL) to create a single-cell suspension. The cells were stained with anti-murine CD11b-FITC (BD Bio-sciences), CD11c-FITC (BD Biosciences), or CD3 FITC (BD Biosciences) antibody, washed in ice-cold PBS, and analyzed using the BD FACS Canto Flow Cytometer.

### Animal immunization with aTEV-based protein and mRNA vaccines

All animal procedures were conducted in accordance with the Guidelines for Animal Experiments and approved by the Animal Experimentation Ethics Committee of Ewha Womans University.

For protein-based vaccination, aluminum hydroxide [Al(OH)₃; Alum] was used as the positive control adjuvant. Six-week-old male BALB/c mice were randomly divided into seven groups (n = 6 per group): control; 0.005, 0.05, 0.5, and 1 µg of RBD encapsulated in aTEVs; and 1 and 5 µg of RBD protein mixed with 300 µg of Alum. Mice were immunized subcutaneously on days 0, 14, and 21, following a previous study with minor modifications [27]. Blood samples were collected on days 7, 21, 30, and 50 for the analysis of RBD-specific IgG and neutralizing antibodies, and stored at –80 °C until use. Spleens were harvested on day 50 for evaluation of cellular immune responses.

To assess short-term storage stability, aTEV-RBD-Pro was lyophilized and stored at 4 °C for 1 week prior to use. Mice were immunized with either fresh or lyophilized aTEV-RBD-Pro on the same three-dose schedule. Serum and spleens were collected as above to assess humoral and cellular immune responses.

For mRNA-based vaccination, BALB/c mice (n = 5 per group) were intramuscularly injected with 1 or 10 µg of aTEV-RBD-mRNA on days 0 and 21. On day 35 (two weeks after the second dose), blood samples were collected for analysis of RBD-specific IgG and neutralizing antibodies, and spleens were harvested for ELISpot and cytokine assays to assess cellular immune responses.

### Measurement of RBD-specific IgG titers

Blood was collected from mice through the retro-orbital plexus. Whole blood was centrifuged at 1,000 × g for 10 min at 4 °C, and serum was collected and stored at –80 °C until use. For RBD-specific IgG quantification, ELISA plates (Corning) were coated with recombinant SARS-CoV-2 RBD protein (2 µg/mL in PBS; Gibco) and incubated overnight at 4 °C. Two columns per plate were coated with 1:1000-diluted goat anti-mouse kappa and lambda light chains (Southern Biotech) as IgG standards. The next day, plates were washed three times with 0.05% Tween-20 in PBS (PBS-T) and blocked with 3% BSA in PBS for 2 h at 37 °C. Serially diluted serum samples or IgG standards were added and incubated for 1 h at 37 °C. After additional washing, goat anti-mouse IgG-HRP (Southern Biotech) was added and incubated for 1 h at room temperature. Signal was developed using TMB substrate (Invitrogen) and stopped with stop solution (R&D Systems). Absorbance was measured at 450 nm using a Synergy H1 Microplate Reader (BioTek), and IgG titers were calculated based on the standard curve.

### SARS-CoV-2 Surrogate virus neutralization test

SARS-CoV-2 Surrogate Virus Neutralization Test (sVNT) kits were obtained from GenScript, Inc. and the tests were performed according to the manufacturer’s instructions. Plasma samples and positive and negative controls were diluted 1:10 and mixed with an equal volume of HRP-conjugated RBD solution. After incubation for 30 min at 37°C, 100 µL each mixture was added to a 96-well plate coated with recombinant ACE2 receptor. After incubation at 37°C for 15 min, the plates were washed four times with wash buffer. Next, 100 µL TMB solution was added and incubated in the dark for 15 min at RT. The reaction was stopped by adding 50 µL Stop Solution and A450 was read using the Synergy H1 Microplate Reader (BioTek). Assay validity was based on the A450 values of the positive and negative controls being in the recommended range. Assuming that the positive and negative controls had acceptable A_450_ values, percentage inhibition was calculated as follows: percentage inhibition = (1 – A_450_ of sample/A_450_ of negative control) × 100. Per-centage inhibition values of < 20% and ≥ 20% were considered negative and positive results, respectively [30].

### ELISPOT assay

Mice were euthanized on day 50. Spleens were harvested and single-splenocyte suspensions were produced. The ELISPOT Kits for murine IFN-γ, TNF-α, and IL-4 were purchased from Cellular Technology Limited (OH, USA). Briefly, splenocytes (8 × 10^5^) were seeded into 96-well plates precoated with the capture antibodies and stimulated in RPMI 1640 medium supplemented with R10 medium (negative control), concanavalin A (2 μg/mL, positive control), or RBD (2 μg/mL). After stimulation, the plates were washed, and spots positive for IFN-γ, TNF-α, and IL-4 were developed according to the manufacturer’s instructions. Spots were counted using the ELISPOT Reader System (Cellular Technology Ltd.); the results are expressed as mean numbers of specific spot-forming cells (SFCs) per 1 × 106 splenocytes.

### Cytokine assay

Splenocytes were harvested for cytokine assays. Splenocyte suspensions from test groups (1 × 10^7^ cells/mL) were plated in 12-well culture plates and incubated with RBD (2 μg/mL) for 48 h at 37°C. Supernatants were harvested and stored at –80°C until analysis of INF-γ, IL-4, TNF-α, and IL 6. Cytokine levels were measured using the Duoset ELISA Kit (R&D Systems) according to the manufacturer’s instructions.

### Statistical analyses

Data are expressed as means ± standard errors of the mean (SEMs) of at least three independent experiments. Statistical analysis was conducted with GraphPad Prism v. 5.0 software. Between-group comparisons were performed using the Mann–Whitney U-test and multiple-group comparisons were done using one way ANOVA.

## Results

### Immunostimulatory effect of EVs derived from LPS-activated THP-1 cells

LPS-stimulated THP-1 cells showed an M1-polarized macrophage phenotype with increased expression of pro-inflammatory cytokines. EVs from M1-polarized, proinflammatory monocytes have an immunostimulatory effect, similar to adjuvants [9, 10]. LPS-activated THP-1 cells showed changes in morphology and increased mRNA levels of pro-inflammatory cytokines such as IL-1β, IL-6, IL-8, TNF-α, and iNOS (Supplementary Fig. 1). The mean diameters of EVs purified from nTEVs and aTEVs, as measured via NTA, were 130.7 ± 1.07 and 141.7 ± 1.41 nm, respectively (Fig. 1A). The nTEVs and aTEVs had similar physical properties. Western blotting showed that both were positive for EV markers (CD9, CD63, CD81, Syntenin 1, Alix, TSG101, and Flotillin-1) and negative for non-EV markers (Calnexin, GM130, and GAPDH) (Fig. 1B and Supplementary Fig.2).).

**Figure 1.**
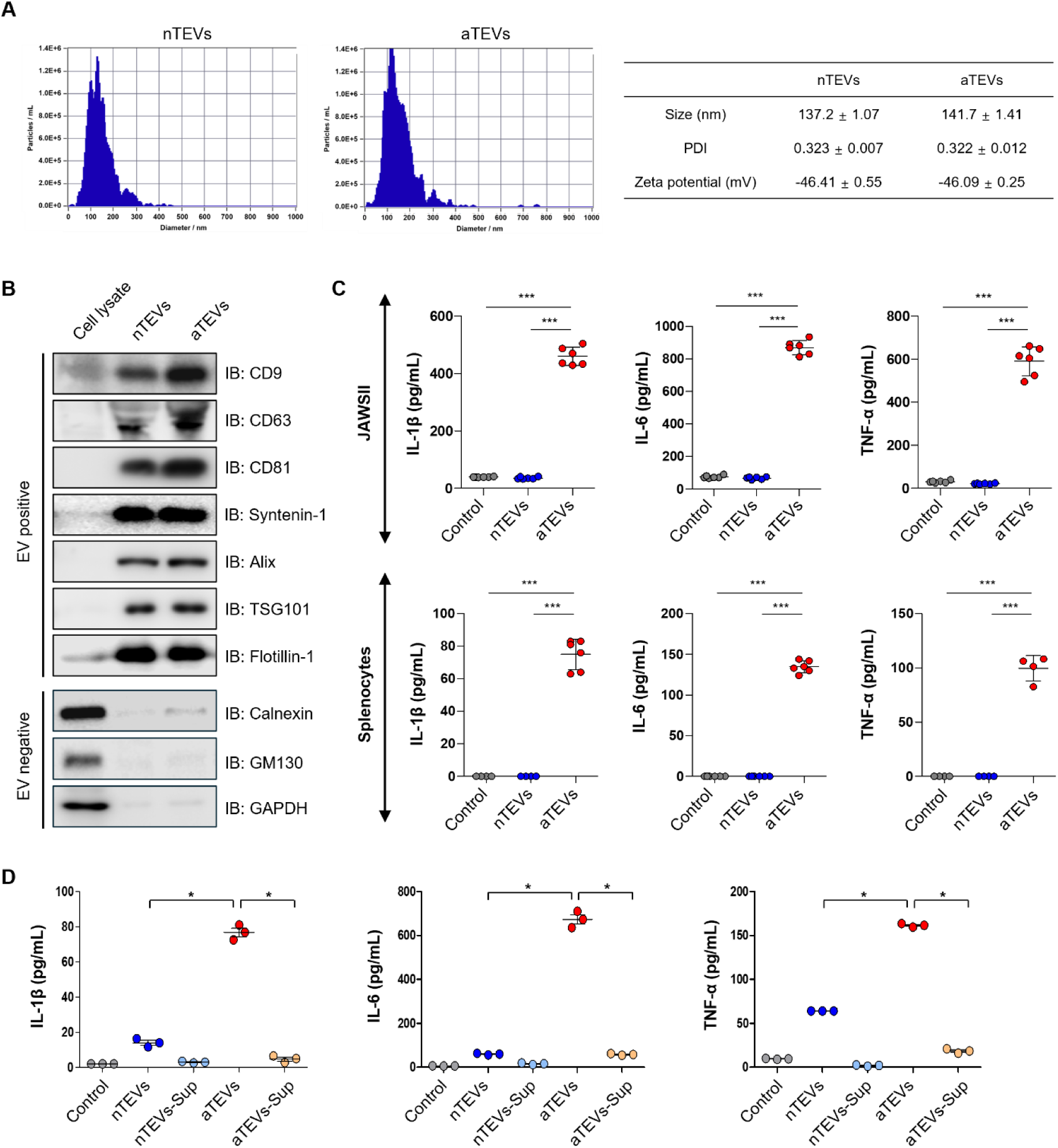
Characterization and immunostimulatory effect of EVs from naïve and LPS-activated THP-1 cells. (A) Size distribution, polydispersity index (PDI), and zeta potential (mV) of nTEVs and aTEVs. (B) Levels of EV markers (CD9, CD63, CD81, Syntenin-1, Alix, TSG101, and Flotillin-1) and non-EV markers (Calnexin, GM130, and GAPDH) as determined by Western blotting. (C) Cytokine levels in JAWSII cells and splenocytes. (D) Cytokine levels in splenocytes treated with TEVs and supernatants of TEVs. nTEVs, naïve THP-1 cell-derived EVs; nTEVs-Sup, supernatant of nTEVs; aTEVs, LPS-activated THP-1 cell-derived EVs; aTEVs-Sup, supernatant of aTEVs; ns, non-significant, *P < 0.05, **P < 0.01, ***P < 0.001.

To evaluate the immunostimulatory activity of purified aTEVs, JAWSII dendritic cells (DCs) or mouse splenocytes were treated with nTEVs or aTEVs and the levels of pro-inflammatory cytokines were analyzed. aTEVs induced high levels of IL-1β, IL-6, and TNF-α in JAWSII cells and splenocytes but nTEVs did not (Fig. 1C). Next, to assess whether the immunostimulatory effect of aTEVs was a result of LPS in the medium, splenocytes were treated with supernatant of EVs, and cytokine production was evaluated. The increased IL-1β, IL-6, and TNF-α levels in cells treated with aTEVs was not a result of residual LPS, confirming the immunostimulatory effect of aTEVs (Fig. 1D). These results suggest that aTEVs from LPS-activated THP-1 cells have potential as an adjuvant because they have an immunostimulatory effect, including triggering the production of pro-inflammatory cytokines.

### In vivo biodistribution and uptake of aTEVs after subcutaneous administration

Delivery of antigens and adjuvant to APCs in lymph nodes is crucial for vaccine effectiveness. aTEVs were labeled with DiR, a lipophilic near-infrared fluorescent cyanine dye, and injected subcutaneously into the right flanks of mice. Their distribution was evaluated using an ex vivo imaging system 24 h after injection. DiR-labeled aTEVs were mainly observed in inguinal and axillary lymph nodes on the ipsilateral but not the contralateral side to the injection site, suggesting that aTEVs migrate directly to the draining lymph nodes rather than via the systemic circulation (Fig. 2A).

**Figure 2.**
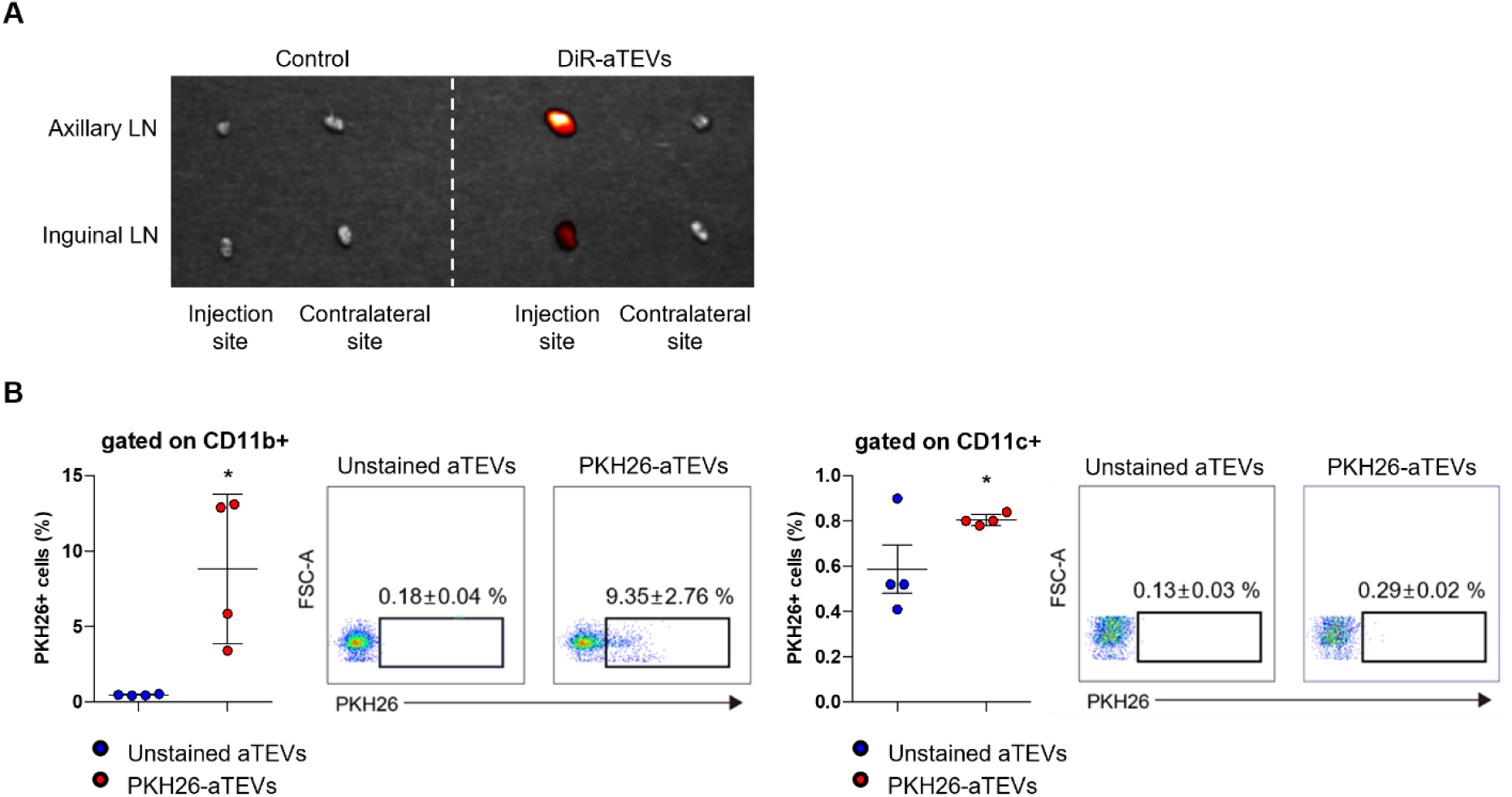
Biodistribution and uptake of aTEVs after subcutaneous injection. DiR-labeled or PKH26-stained aTEVs were subcutaneously injected into the right flank of mice and the lymph nodes were removed 24 h later. (A) Representative ex vivo images of the axillary and inguinal lymph nodes of a mouse injected with DiR-labeled aTEVs. (B) Representative FACS data of the cell populations in lymph nodes that took up PKH26-stained aTEVs. *P < 0.05.

Next, flow cytometry was performed to identify the lymph node cells that take up aTEVs. Cells from inguinal and axillary lymph nodes were stained with antibodies for the macrophage marker CD11b, DC marker CD11c, and the T-cell marker CD3. aTEVs were taken up mainly by macrophages in the lymph nodes; <1% were taken up by DCs and T cells (Fig. 2 and Supplementary Fig. 3). These results suggest that subcutaneously injected aTEVs preferentially migrate to draining lymph nodes, where they are mainly taken up by macrophages. However, systemic distribution to non-lymphoid organs cannot be excluded at this stage.

### High-efficiency encapsulation and delivery of SARS-CoV-2 RBD protein and mRNA in aTEVs by SW

The high biocompatibility, low toxicity, nanoscale size, and cargo-loading ability of extracellular vesicles (EVs) make them attractive carriers for antigen delivery [26, 27]. To develop a flexible EV-based vaccine platform, we used shock wave (SW) treatment to encapsulate either SARS-CoV-2 receptor-binding domain (RBD) protein or RBD mRNA into immunostimulatory EVs (aTEVs) derived from LPS-stimulated THP-1 monocytes. First, we evaluated the effect of SW treatment on the physical properties of EVs using nanoparticle tracking analysis (NTA), dynamic light scattering (DLS), zeta potential analysis, and cryo-electron microscopy (cryo-EM). The size, polydispersity index (PDI), and zeta potential of aTEVs were not significantly altered by SW (Fig. 3A), and cryo-EM imaging confirmed preservation of the spheroid vesicle morphology (Fig. 3B). When 1000 pulses of SW at 0.3 mJ/mm² were applied in the presence of RBD protein, the encapsulation efficiency reached 68.94 ± 2.58% without evidence of vesicle damage (Fig. 3C, D). To confirm delivery, aTEV-RBD-Pro was incubated with HEK-293T cells, and intracellular RBD protein expression was assessed by Western blotting. Only the SW-encapsulated group showed detectable RBD signal, whereas cells treated with aTEVs mixed with free RBD protein did not (Fig. 3E), indicating that encapsulation is essential for cellular de-livery.

**Figure 3.**
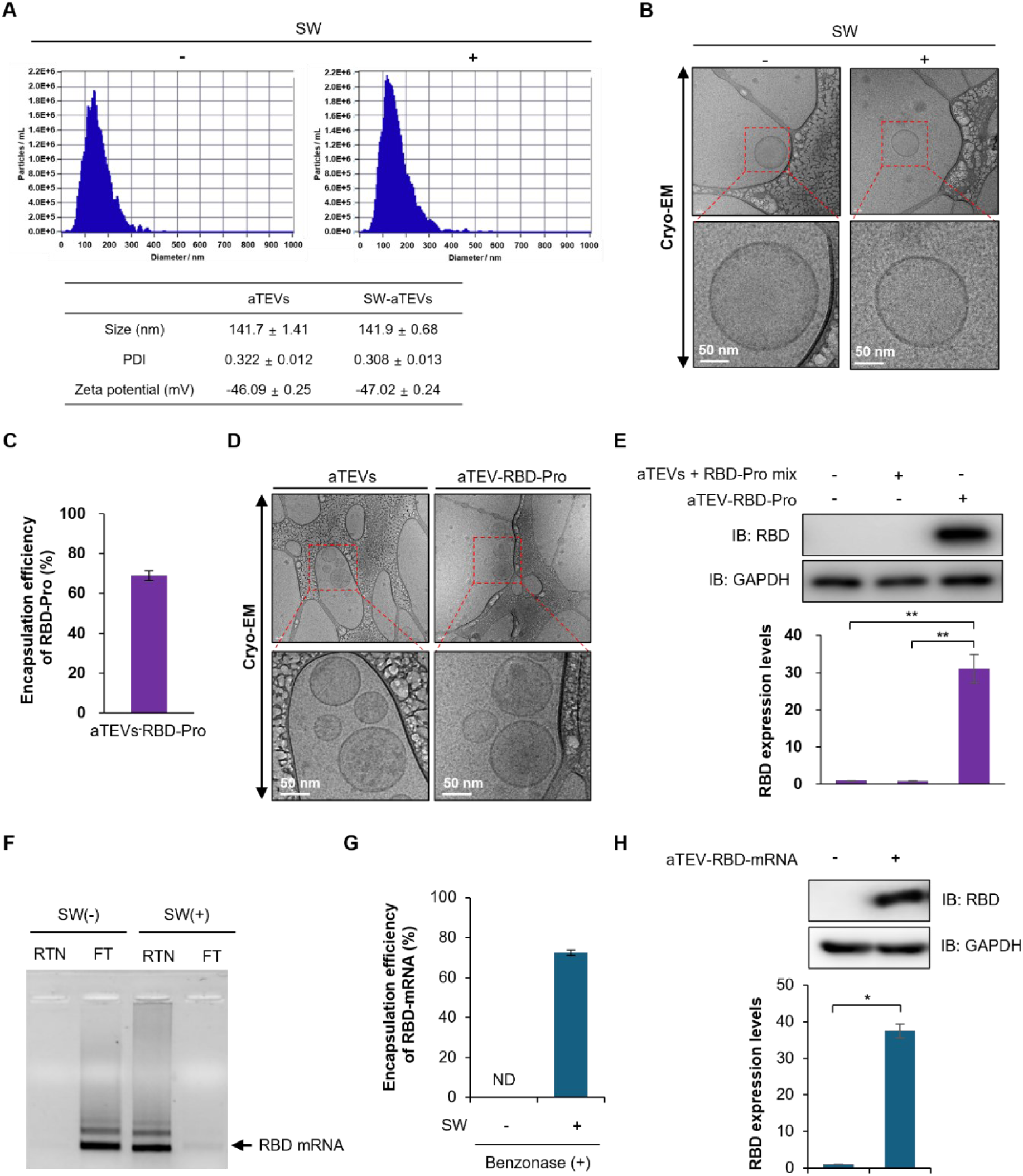
Encapsulation and delivery of SARS-CoV-2 RBD protein and mRNA into aTEVs using SW. (A) Size distribution, polydispersity index (PDI), and zeta potential (mV) of aTEVs before and after SW treatment, showing no significant change in physicochemical properties. (B) Representative cryo-EM images of aTEVs before and after SW application (scale bars, 50 nm), confirming preserved vesicle morphology. (C) Quantification of RBD protein encapsulation efficiency in aTEVs following SW treatment (∼69%). Encapsulation efficiency of SARS-CoV-2 RBD protein into aTEVs using shock wave treatment. This high efficiency (∼69%) supports the use of SW as a non-destructive, effective method for post-loading of protein antigens into EVs, which contrasts with lower efficiency rates often reported with electroporation or sonication (D) Cryo-EM images of aTEVs after RBD protein encapsulation, demonstrating vesicle integrity. (E) Western blot analysis of RBD protein expression in HEK-293T cells treated with either free RBD + aTEVs (1 μg) or RBD protein–loaded aTEVs (aTEV-RBD-Pro, equivalent to 1 μg RBD). *P < 0.05. (F) RBD mRNA encapsulation into aTEVs with or without shock wave (SW) treatment, assessed using 100 kDa molecular weight cut-off filtration and agarose gel electrophoresis. aTEVs were incubated with RBD mRNA and either treated with SW or left untreated. In the absence of SW, mRNA was detected only in the filtrate (FT), indicating lack of EV association. In contrast, SW-treated samples showed strong mRNA retention in the EV-containing retentate (RT), with only a barely detectable signal in the filtrate, con-firming that SW is required for efficient mRNA loading into EVs. (G) Quantification of internalized RBD mRNA in aTEVs after benzonase treatment. aTEVs treated with or without SW were exposed to benzonase to eliminate surface-bound RNA, followed by RT-qPCR. Approximately 85% of total RNA signal was retained in the SW group, confirming effective mRNA encapsulation. (H) Western blot detection of intracellular RBD protein in HEK-293T cells treated with aTEV-RBD-mRNA. A clear RBD band was detected, demonstrating functional delivery and translation of the mRNA cargo.

We next applied the SWEET method to load RBD mRNA into aTEVs under the same SW conditions. To evaluate the effect of SW treatment on mRNA loading, aTEVs were mixed with RBD mRNA and either subjected to SW or left untreated. Agarose gel electrophoresis revealed that in the absence of SW, RBD mRNA was detected primarily in the filtrate after 100 kDa ultrafiltration, indicating that it remained free in solution and was not associated with EVs. In contrast, SW-treated samples showed strong mRNA retention in the EV-containing retentate, with only a barely detectable signal in the filtrate, demonstrating that SW treatment enables effective incorporation of mRNA into EVs (Fig. 3F) Furthermore, to distinguish internalized mRNA from surface-bound RNA, aTEVs were treated with Benzonase to degrade unprotected nucleic acids. RT-qPCR quantification showed that approximately 72% of the RBD mRNA signal was retained in SW-treated aTEVs, whereas untreated controls exhibited substantially lower protected RNA levels (Fig. 3G). These data confirm that mRNA was successfully internalized into EVs by SW. Finally, we evaluated the functional delivery of aTEV-RBD-mRNA. Western blotting confirmed the presence of intracellular RBD protein in HEK-293T cells treated with mRNA-loaded aTEVs, indicating that the mRNA cargo was successfully delivered and translated into protein (Fig. 3H). To further validate the general applicability of SWEET-mediated mRNA loading, we applied the same method to deliver EGFP mRNA using milk-derived EVs. When HCT116 cells were treated with milk EVs containing EGFP mRNA, robust GFP expression was observed in over 90% of cells by fluorescence microscopy and flow cytometry (Supplementary Fig. 4), demonstrating that SWEET enables functional mRNA delivery across different EV types. These results demonstrate that the SWEET platform enables efficient, quantifiable encapsulation and intracellular delivery of both protein and mRNA antigens using the same EV-based system.

### Humoral immune responses induced by aTEV-RBD-Pro and aTEV-RBD-mRNA

To evaluate the potential of aTEV-RBD-Pro as a protein-based vaccine, mice were immunized via subcutaneous (SC) injection of 0.005–1 µg of RBD protein encapsulated in aTEVs on days 0, 14, and 21. As a positive control, 1 or 5 µg of RBD protein formulated with alum adjuvant (alum) was administered subcutaneously using the same schedule. Serum samples were collected on days 7, 21, 30, and 50 for ELISA-based quantification of RBD-specific IgG levels. Mice immunized with ≥ 0.1 µg aTEV-RBD-Pro showed dose-dependent induction of RBD-specific IgG as early as day 7, and this response was sustained through day 50. In contrast, mice that received alum-adjuvanted RBD protein showed minimal IgG induction at day 7, although robust responses were observed from day 21 onward. Among mice receiving 1 µg RBD, the aTEV-RBD-Pro group exhibited IgG levels comparable to those in the alum-adjuvanted RBD-Pro (Adj+RBD-Pro) groups (Fig. 4A). To assess functional antibody responses, neutralizing activity was measured at day 50 using a surrogate virus neutralization test (sVNT). Mice immunized with 0.05 µg or more of aTEV-RBD-Pro showed inhibition rates exceeding the 20% positive cutoff, with 1 µg aTEV-RBD-Pro inducing 87 ± 2.21% inhibition, comparable to the 95.1 ± 0.7% observed in the Adj+RBD-Pro control (Fig. 4B). Statistical analysis indicated a significant dose-dependent increase in both IgG titers and neutralizing activity (P < 0.05 to **P < 0.001).

**Figure 4.**
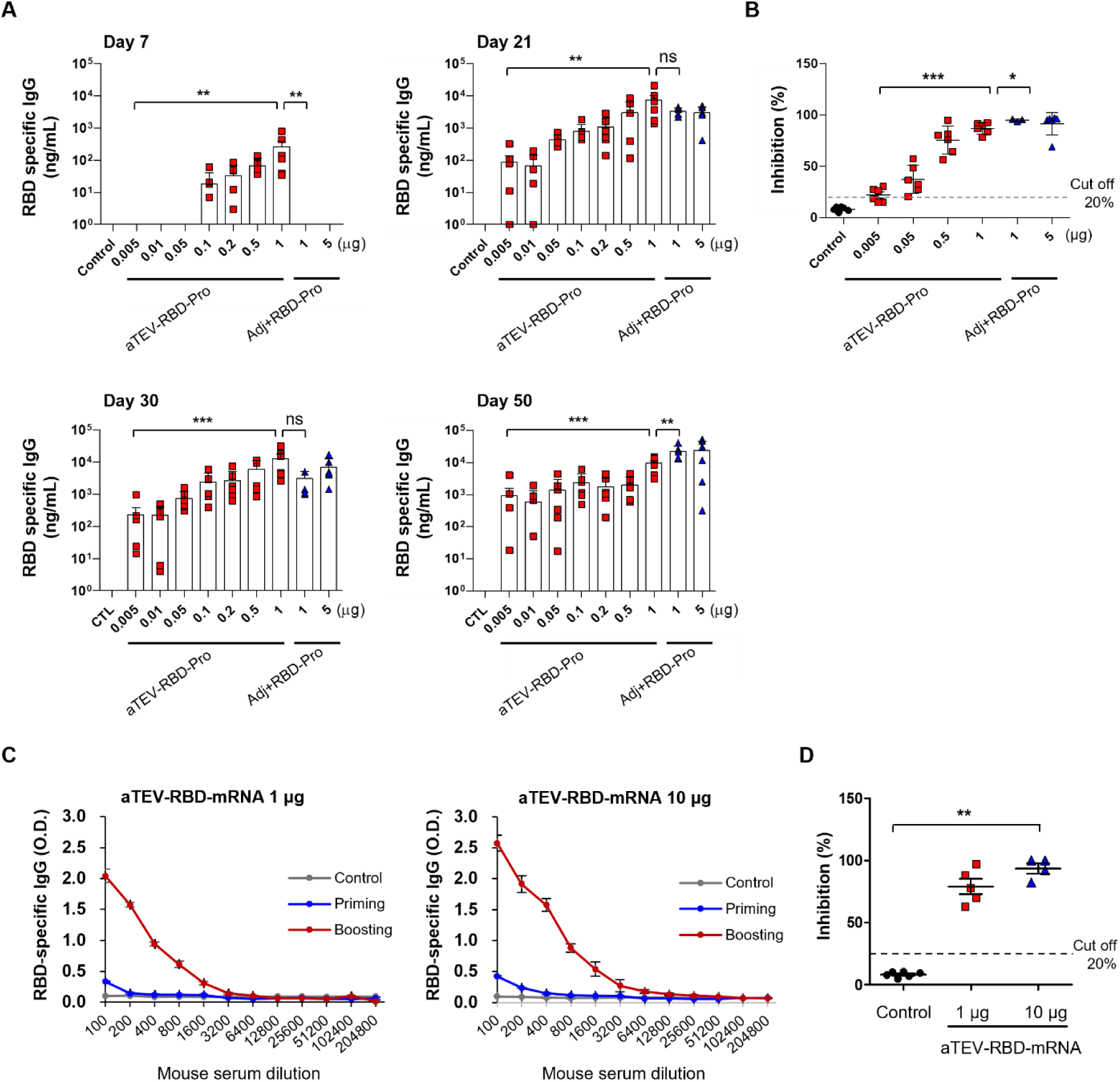
Humoral immune responses induced by aTEV-RBD-Pro and aTEV-RBD-mRNA vaccination. (A, B) BALB/c mice were immunized with aTEV-RBD-Pro (0.005–1 µg, subcutaneously on days 0, 14, and 21) or with alum-adjuvanted RBD protein (Adj+RBD-Pro, 1 or 5 µg) as a control. (A) RBD-specific IgG titers in serum at days 7, 21, 30, and 50, measured by ELISA. (B) Neutralizing antibody activity in serum at day 50, assessed using the SARS-CoV-2 surrogate virus neutralization test (sVNT). (C, D) BALB/c mice were immunized with aT-EV-RBD-mRNA (1 or 10 µg, intramuscularly on days 0 and 21). (C) RBD-specific IgG titers in serum at day 35 (14 days after the second dose), measured by ELISA. (D) Neutralizing antibody activity in serum at day 35, assessed by sVNT. ns, non-significant; *P < 0.05, **P < 0.01, ***P < 0.001.

Next, we evaluated humoral responses induced by aTEV-RBD-mRNA, delivered via intramuscular (IM) injection. Mice received 1 or 10 µg of mRNA-encapsulated aTEVs on days 0 and 21. Serum was collected on day 35 (14 days after the second injection) for antibody analysis. ELISA revealed a clear dose-dependent increase in RBD-specific IgG titers, with significantly higher levels in the 10 µg group (Fig. 4C). Consistently, sVNT analysis showed that both 1 µg and 10 µg groups generated neutralizing antibodies capable of inhibiting the RBD–ACE2 interaction, with stronger activity observed in the higher-dose group (Fig. 4D). These results confirm that both protein and mRNA formulations of aTEV-based vaccines elicit potent and functional humoral immune responses, even in the absence of external adjuvants.

### Cellular immune responses induced by aTEV-RBD-Pro and aTEV-RBD-mRNA

To evaluate antigen-specific cellular immune responses following immunization, spleens were harvested and splenocytes were analyzed using ELISpot assays and cytokine profiling. Splenocytes from mice immunized with aTEV-RBD-Pro were stimulated in vitro with recombinant RBD protein. The ELISpot assay revealed robust secretion of IFN-γ, IL-4, and TNF-α in aTEV-RBD-Pro–immunized mice, with clear dose-dependent increases (Fig. 5A). The highest cellular responses were observed in the 1 µg group, whereas the alum-adjuvanted RBD (Adj+RBD-Pro) groups elicited only minimal or background-level cytokine responses. Similarly, cytokine quantification from splenocyte culture supernatants showed increased production of IFN-γ, IL-4, TNF-α, IL-2, and IL-6 in aTEV-RBD-Pro–treated mice in a dose-dependent manner (Fig. 5B). These results indicate that EV-encapsulated RBD protein strongly activates both Th1- and Th2-associated cytokine responses without the need for external adjuvants.

**Figure 5.**
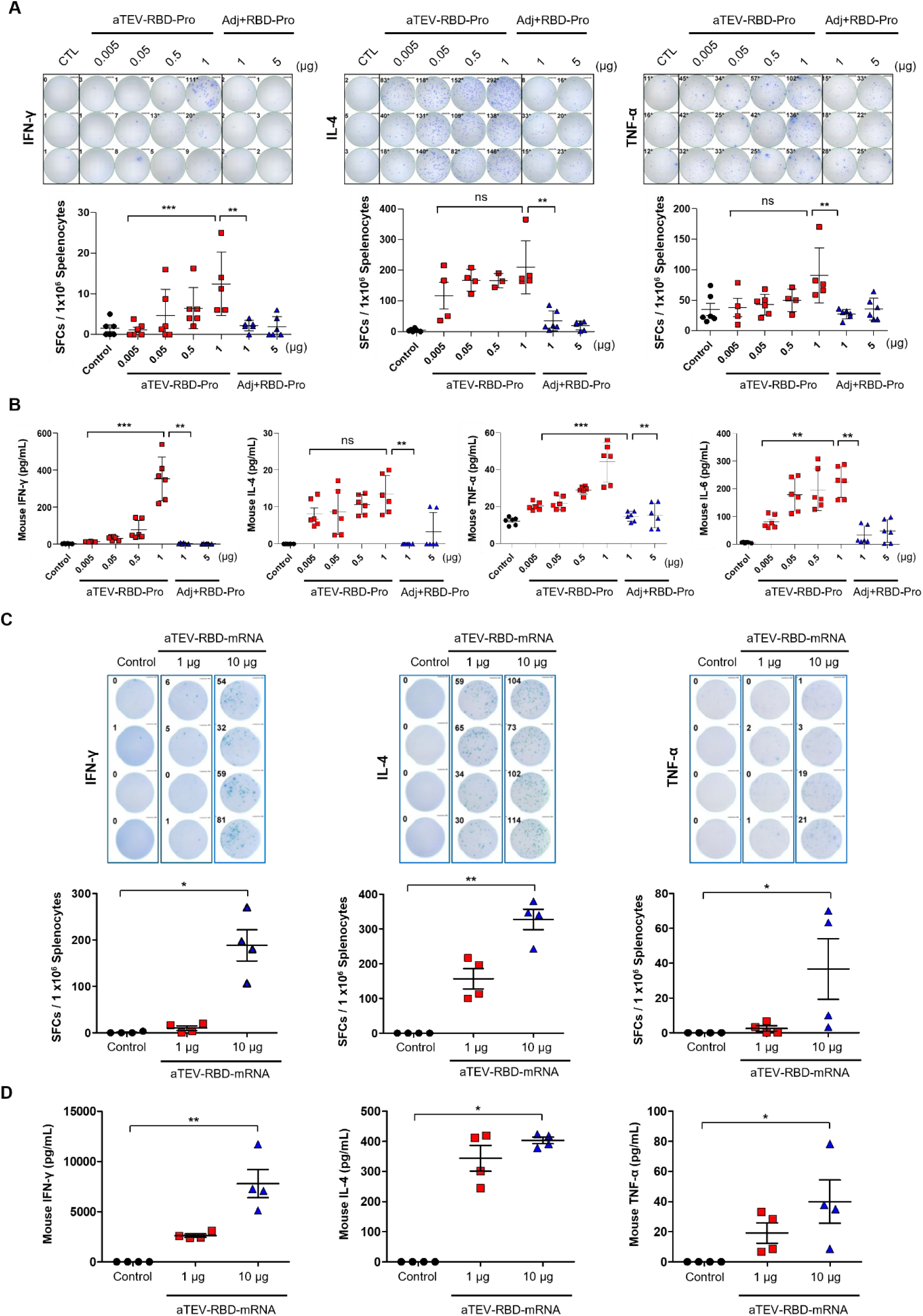
Induction of cellular immune responses in mice immunized with aTEV-based vaccines. (A, B) ELISpot and cytokine assays of splenocytes isolated from mice immunized with aTEV-RBD-Pro or alum-adjuvanted RBD (Adj+RBD-Pro). (A) ELISpot assay quantifying IFN-γ, IL-4, and TNF-α secretion in antigen-restimulated splenocytes. Spot-forming units (SFUs) were calculated per 10⁶ cells. Dose-dependent in-creases in cytokine-producing cells were observed in the aTEV-RBD-Pro groups. (B) Cytokine concentrations (IFN-γ, IL-4, TNF-α, IL-2, and IL-6) in culture supernatants measured by multiplex assay. (C, D) Cellular immune responses in mice immunized with aTEV-RBD-mRNA. (C) ELISpot analysis of IFN-γ, IL-4, and TNF-α production from antigen-restimulated splenocytes. (D) Quantification of IFN-γ, IL-4, and TNF-α in culture supernatants by multiplex assay.

We also evaluated whether antigen loading into EVs via SW is necessary for cellular immunity. Mice immunized with a simple mixture of aTEVs and RBD protein (without SW treatment) produced lower levels of RBD-specific IgG compared to those immunized with aTEV-RBD-Pro or Adj+RBD-Pro (Supplementary Fig. 5A). Neutralizing antibody production was minimal in this group (Supplementary Fig. 5B), although a modest RBD-specific cellular response was observed (Supplementary Fig. 5C). These results suggest that the aTEVs themselves possess inherent immunostimulatory properties, but efficient antigen delivery via SW-mediated encapsulation is essential for eliciting optimal humoral and cellular responses.

For the mRNA vaccine, BalB/C mice were immunized intramuscularly with 1 µg or 10 µg of aTEV-RBD-mRNA on days 0 and 21. On day 35, ELISpot analysis showed significantly increased numbers of IFN-γ–, IL-4–, and TNF-α–producing splenocytes in a dose-dependent manner (Fig. 5C). Multiplex cytokine analysis of splenocyte supernatants confirmed increased levels of these cytokines, particularly in the 10 µg group (Fig. 5D). These results confirm that aTEV-RBD-mRNA also effectively induces antigen-specific cellular immunity, with comparable Th1 and Th2 cytokine profiles to the protein-based vaccine, despite differences in antigen format, delivery route, and schedule.

Together, these findings demonstrate that both aTEV-RBD-Pro and aTEV-RBD-mRNA vaccines induce functional cellular immune responses without the need for traditional adjuvants, and highlight the essential role of SWEET-mediated antigen encapsulation for effective immunization.

### Storage stability of lyophilized aTEV-RBD-Pro

To evaluate their storage stability, aTEV-RBD-Pro was lyophilized and stored at 4°C for 1 week and their induction of IgG and neutralizing antibodies was evaluated. Lyophilized aTEV-RBD-Pro induced similar levels of RBD-specific IgG and neutralizing antibodies as fresh aTEV-RBD-Pro (Fig. 6A and 6B).

**Figure 6.**
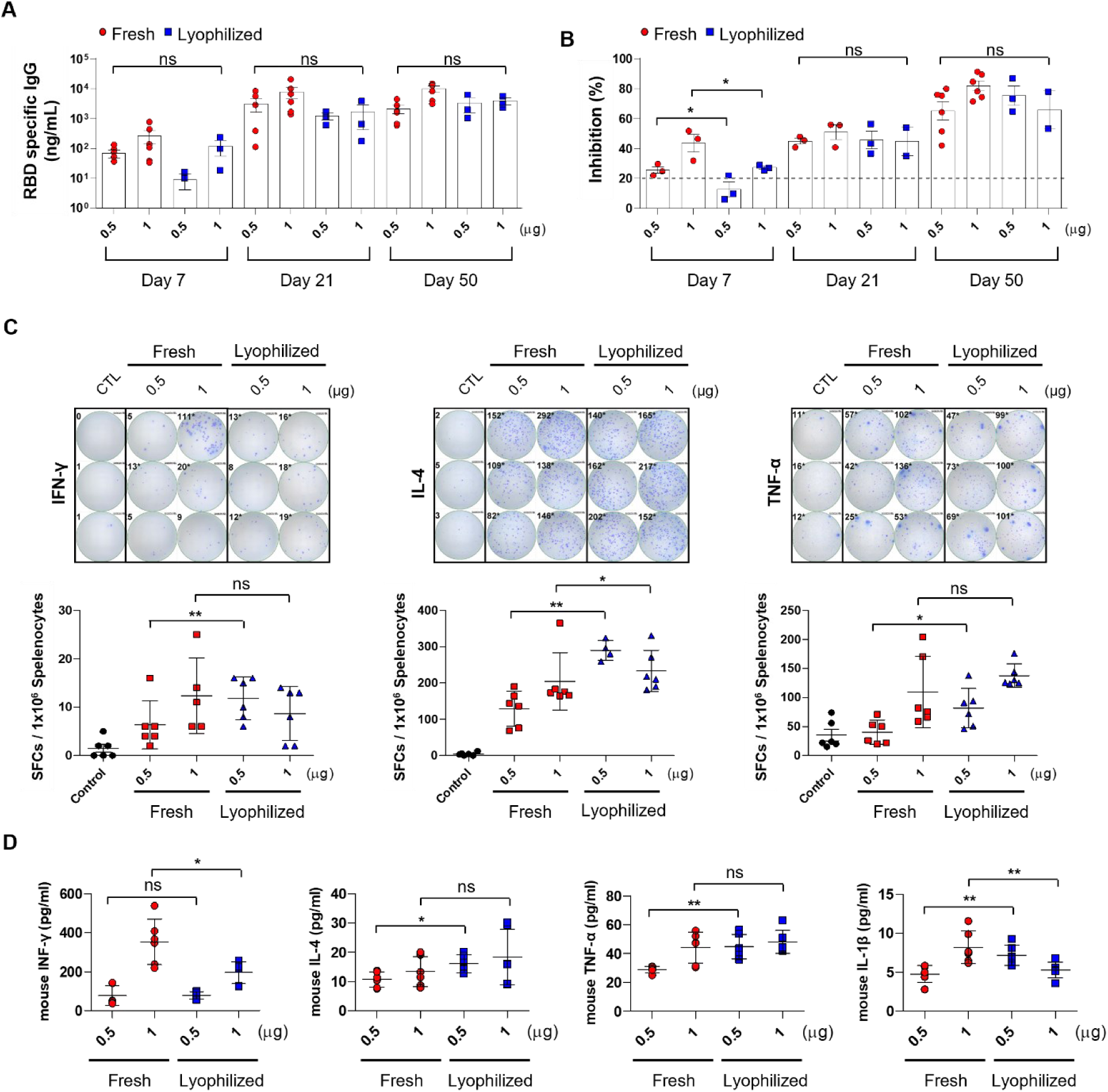
Storage stability of lyophilized aTEV-RBD-Pro. BALB/c mice (n = 6/group) were immunized subcutaneously on days 0, 14, and 21 with either freshly prepared or lyophilized aTEV-RBD-Pro (containing 0.5 or 1 µg of encapsulated RBD protein), with lyophilized samples stored at 4 °C for 7 days prior to use. (A) RBD-specific IgG levels in serum measured by ELISA at days 7, 21, and 50. (B) Neutralizing antibody activity in serum at days 7, 21, and 50, determined using the SARS-CoV-2 surrogate virus neutralization test (sVNT). (C) ELISpot assay to quantify RBD-specific IFN-γ–, IL-4–, and TNF-α–producing cells in splenocytes. (D) Cytokine concentrations in splenocyte culture supernatants, measured by multiplex bead-based assay. ns, non-significant; *P < 0.05, **P < 0.01.

ELISpot and cytokine assays for IFN-γ, IL-4, and TNF-α indicated induction of an RBD-specific cellular immune response in mice immunized with lyophilized aTEV-RBD-Pro (Fig. 6C and 6D). These findings suggest that EV-based vaccines would have good storage stability, obviating the need for cold-chain distribution.

## Discussion

In this study, we developed a modular extracellular vesicle (EV)-based vaccine platform by leveraging the immunostimulatory properties of activated monocyte-derived EVs and applying shock wave (SW)-mediated post-loading. EVs derived from lipopolysaccharide (LPS)-stimulated THP-1 monocytes exhibited inherent adjuvant-like activity and were used to deliver both SARS-CoV-2 receptor-binding domain (RBD) protein and RBD-encoding mRNA. Notably, both cargo formats elicited robust antigen-specific humoral and cellular immune responses in mice without the need for additional adjuvants, highlighting the dual functionality of these engineered EVs (aTEVs) as both delivery vehicles and immunostimulatory agents (Figs. 4–6 and Supplementary Fig. 5).

This strategy builds upon and advances previous EV-based vaccine approaches. Exosome vaccines carrying viral antigens have been shown to induce cytotoxic T lymphocyte (CTL) responses [13], and exosomes derived from M1 macrophages enhanced vaccine efficacy by promoting inflammation in draining lymph nodes [9]. More recently, Luo et al. reported an EV vaccine platform that simultaneously delivers mRNA and protein. However, their system relies on lentiviral transduction of 293F cells to overexpress the antigen, allowing endogenous incorporation of mRNA and protein into EVs [28]. This approach is inherently limited by its dependence on producer cell biology, the inability to quantify true encapsulation efficiency, and the lack of direct evidence for functional delivery. Specifically, the amount of antigen associated with EVs was inferred from relative mRNA Ct values and estimated protein proportions, and most of the expressed antigen remained outside the EV fraction. In contrast, our approach enables precise post-loading of pre-purified antigens into EVs using SWEET technology. We achieved high-efficiency encapsulation of RBD protein (∼69%) and confirmed RBD mRNA presence via electrophoresis after SW-mediated loading. Functional delivery was demonstrated by intracellular detection of RBD protein following treatment with aTEV-RBD-mRNA, supporting successful translation of EV-delivered mRNA. This level of control and validation is a key advantage over cell-based passive loading strategies. The superior encapsulation efficiency of the SWEET platform was validated in a subsequent study published in 2024, where shock wave–based EV loading outperformed electroporation, sonication, and lipofection in terms of RNA delivery efficiency [25].

Importantly, the immunostimulatory effects of aTEVs were not due to residual LPS, as confirmed by endotoxin assays (Supplementary Figure 6). Instead, pro-inflammatory cytokines and surface molecules intrinsic to LPS-activated monocytes are likely contributors to immune potentiation.

Cryo-EM and particle analysis further confirmed that SW treatment preserved vesicle morphology, supporting its utility as a non-destructive and scalable loading method. EVs have been widely reported to exhibit excellent biocompatibility and negligible systemic toxicity, even when distributed beyond their intended target tissues. In our study, no adverse effects were observed in vaccinated animals, supporting the inherent safety of the aTEV-based formulation. Although our biodistribution analysis showed selective ac-cumulation of aTEVs in draining lymph nodes after SC injection, we acknowledge that EVs may systemically distribute to major organs, particularly the liver and spleen, in other administration contexts. EV uptake in the liver—primarily by Kupffer cells and occasionally by hepatocytes—is a well-documented phenomenon in EV research. While we did not observe liver toxicity or adverse effects in this study, future work will include comprehensive organ-level biodistribution and histopathological analysis to further characterize off-target uptake and ensure translational robustness. Functionally, both aTEV-RBD-Pro and aTEV-RBD-mRNA induced potent immune responses in vivo, comparable or superior to those elicited by conventional adjuvanted vaccines. Lyophilization preserved vaccine activity, eliminating the need for cold-chain logistics—an essential consideration for global deployment.

To evaluate immune activation, we analyzed key Th1/Th2 cytokines including IFN-γ, IL-4, TNF-α, and IL-6. These markers are commonly used in vaccine studies and are particularly relevant to SARS-CoV-2, where IFN-γ and IL-6 have been linked to protective immunity [2, 29]. While other cytokines such as IL-12, IL-17, and IL-23 are also important in viral clearance—especially at mucosal sites—we focused on classical systemic cytokines such as IFN-γ, IL-4, TNF-α, and IL-6 in this initial characterization due to sample limitations. The selection of specific cytokines also varied across experiments depending on the immunological objective and biological context. IL-1β was assessed in in vitro assays (Figure 1) to evaluate innate inflammatory responses, particularly inflammasome activation. In contrast, IL-6, IFN-γ, TNF-α, and IL-4 were measured in in vivo studies (Figures 5–6) to characterize adaptive immune polarization and effector function. This design reflects the distinct mechanistic goals of each assay. Future studies will broaden the cytokine panel to include additional pathways to fully elucidate the immune landscape induced by aTEV-RBD vaccination.

While both aTEV-RBD-Pro and aTEV-RBD-mRNA vaccines successfully elicited robust humoral immune responses, the experimental conditions used for each format differed in several aspects that should be considered when interpreting and comparing their immunogenicity. The protein-based vaccine was administered subcutaneously with three doses at two-week intervals, whereas the mRNA-based vaccine was delivered intramuscularly with only two doses. Furthermore, the total administered antigen dose differed: up to 1 µg of recombinant RBD protein was used in the aTEV-RBD-Pro group, compared to 1 or 10 µg of mRNA in the aTEV-RBD-mRNA group, which relies on in vivo translation for antigen expression. These differences in delivery route, dosage, and antigen processing pathways could all influence the magnitude and kinetics of the immune response. Subcutaneous injection offers rapid lymphatic drainage to draining lymph nodes, potentially enhancing early T and B cell priming, while intramuscular injection may result in slower antigen exposure. The three-dose schedule of the protein vaccine may also explain the more sustained antibody levels observed through day 50, compared to the two-dose mRNA schedule assessed at day 35. Given the lack of matched conditions across groups, direct comparisons between the two vaccine formats must be made with caution. Nevertheless, both formulations demonstrated dose-dependent IgG responses and functional neutralizing activity without the need for exogenous adjuvants. Future studies using harmonized protocols—including equivalent routes, antigen doses, and timing— will be required to directly compare the immunogenicity and durability of protein- and mRNA-based aTEV vaccines.

One of the key strengths of our platform is its capacity to accommodate either protein or mRNA vaccines using the same EV engineering and loading protocol. This versatility is critical for addressing pathogen-specific or population-specific needs. For instance, mRNA vaccines may offer rapid development and stronger CD8⁺ T cell activation, which are essential for emerging or mutating viruses, while protein-based vaccines may provide better safety profiles and antibody induction, particularly in pediatric or immunocompromised populations. Our system enables this format flexibility without compromising delivery efficacy or immune activation. Because SWEET-based post-loading decouples cargo incorporation from cell-derived production, the platform may also support co-loading of multiple antigen types—including mRNAs or proteins targeting different variants or even pathogens. This opens possibilities for multivalent COVID-19 vaccines or combination vaccines (e.g., SARS-CoV-2 + influenza) using a single EV formulation. Such modularity would be difficult to achieve in cell-dependent systems and offers a powerful tool for rapid response to future outbreaks. The SWEET platform is composed entirely of clinically familiar components: extracellular vesicles, which are already under investigation in human therapies (e.g., MSC-EVs), and acoustic shock waves, which are FDA-approved for multiple non-invasive medical uses. This biocompatible, cell-free, and non-viral formulation may facilitate regulatory approval and clinical scalability. More-over, EVs loaded with defined antigen doses can be readily characterized, which aligns with emerging guidelines for EV-based therapeutics and vaccines.

While our results clearly establish the feasibility and potency of the SWEET-based aTEV platform, several limitations remain. First, we did not perform a live virus challenge to directly measure protective efficacy. Second, the current study focused on a single antigen (RBD) and intramuscular/subcutaneous administration routes. Future studies should extend this platform to multivalent or pan-coronavirus designs, assess long-term immune memory, and explore mucosal delivery. In addition, protease-based confirmation of protein encapsulation will help further distinguish between encapsulated and surface-adsorbed cargoes.

In summary, this work introduces a versatile EV-based vaccine platform that combines innate immune stimulation, post-loading flexibility, and high delivery efficiency for both protein and mRNA antigens. Unlike prior EV vaccine strategies that rely on endogenous cargo expression, our SWEET platform allows dose-defined, quantifiable, and functionally verified antigen delivery. To our knowledge, this is the first report demonstrating that a single engineered EV system can independently deliver both protein and mRNA vaccines with confirmed intracellular expression. These findings provide a strong foundation for further development of EV-based vaccines targeting a broad range of infectious diseases and therapeutic indications.

## Institutional Review Board Statement

The study was conducted according to the guidelines of the Declaration of Helsinki, and approved by the Animal Experimentation Ethics Committee of Ewha Womans University (protocol code EUMC-IACUC-22-006, date of approval: 31th, May, 2022).

## Informed Consent Statement

Not applicable.

## Data Availability Statement

Data is contained within the article or supplementary material.

