## Supplemental data for "A single extracellular vesicle-based platform supporting both RBD protein and mRNA vaccination against SARS-CoV-2"

### Supplemantal data

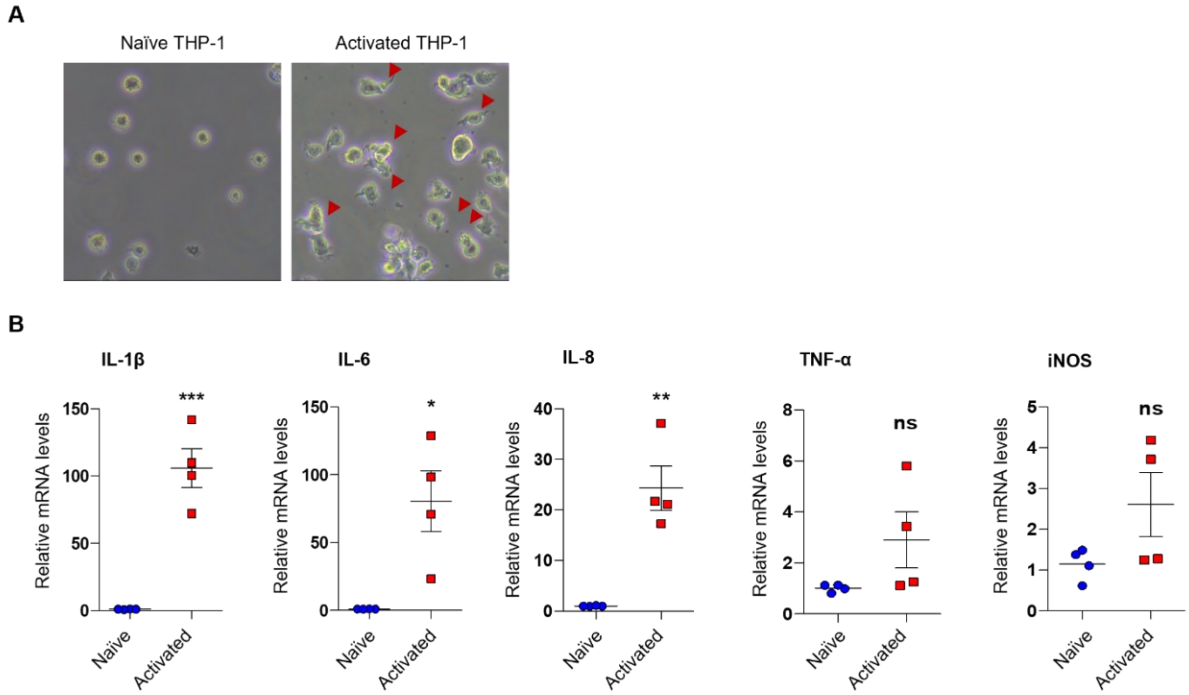


**Supplementary Figure 1.** Phenotype of M1-polarized macrophages in LPS-activated THP-1 cells. (A) Morphology of naïve and LPS-activated THP-1 cells. THP-1 cells were stimulated with 100 ng/mL LPS for 6 h. (B) Relative mRNA levels of pro-inflammatory cytokines in LPS-activated THP-1 cells. ns, non-significant; *P < 0.05, **P < 0.01, ***P < 0.001.

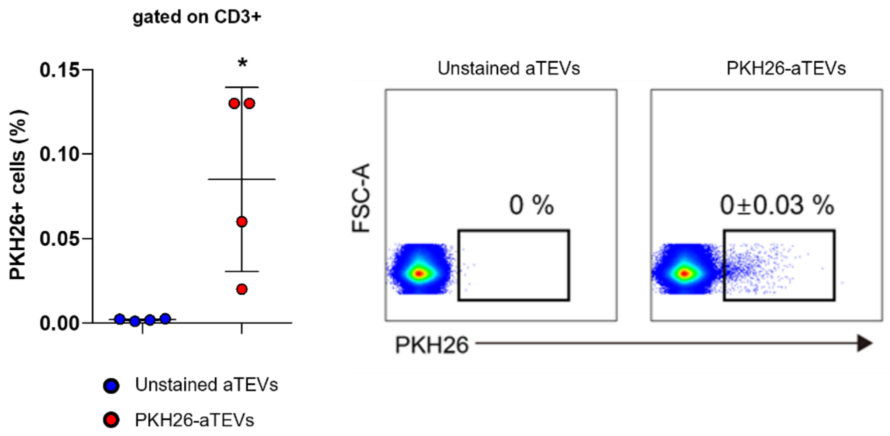


**Supplementary Figure 2**. Uptake of aTEVs by T cells in lymph nodes. PKH26-stained aTEVs were injected subcutaneously into the right flanks of mice. Representative FACS data showing uptake of PKH26-stained aTEVs by CD3+ T cells in lymph nodes. *$P < 0.05$.

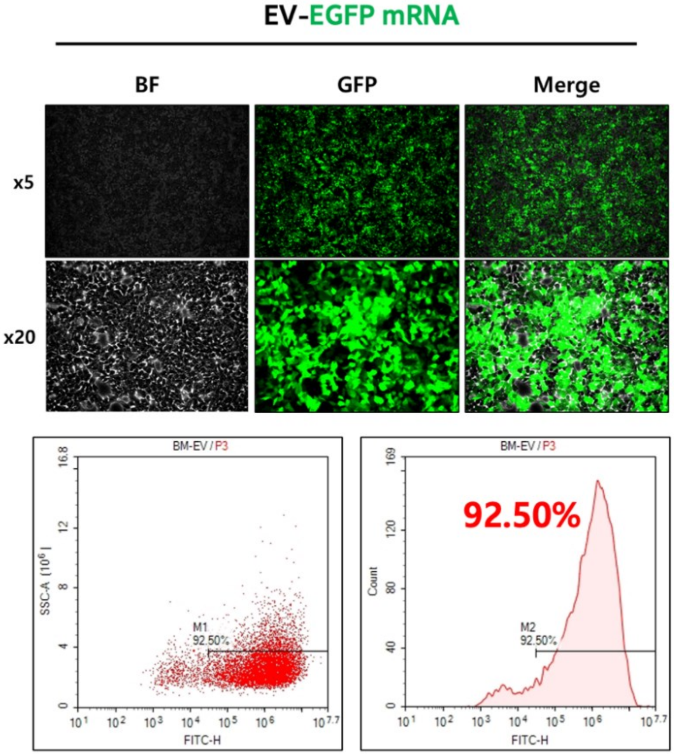


**Supplementary Figure 3**. Expression of EGFP mRNA encapsulated milk EV. The EGFP mRNA was encapsulated into milk EVs using acoustic shockwave and delivered to HCT116 cells. FACS and fluorescence microscopy analyses showed that fluorescence was detected in more than 90% of the cells.

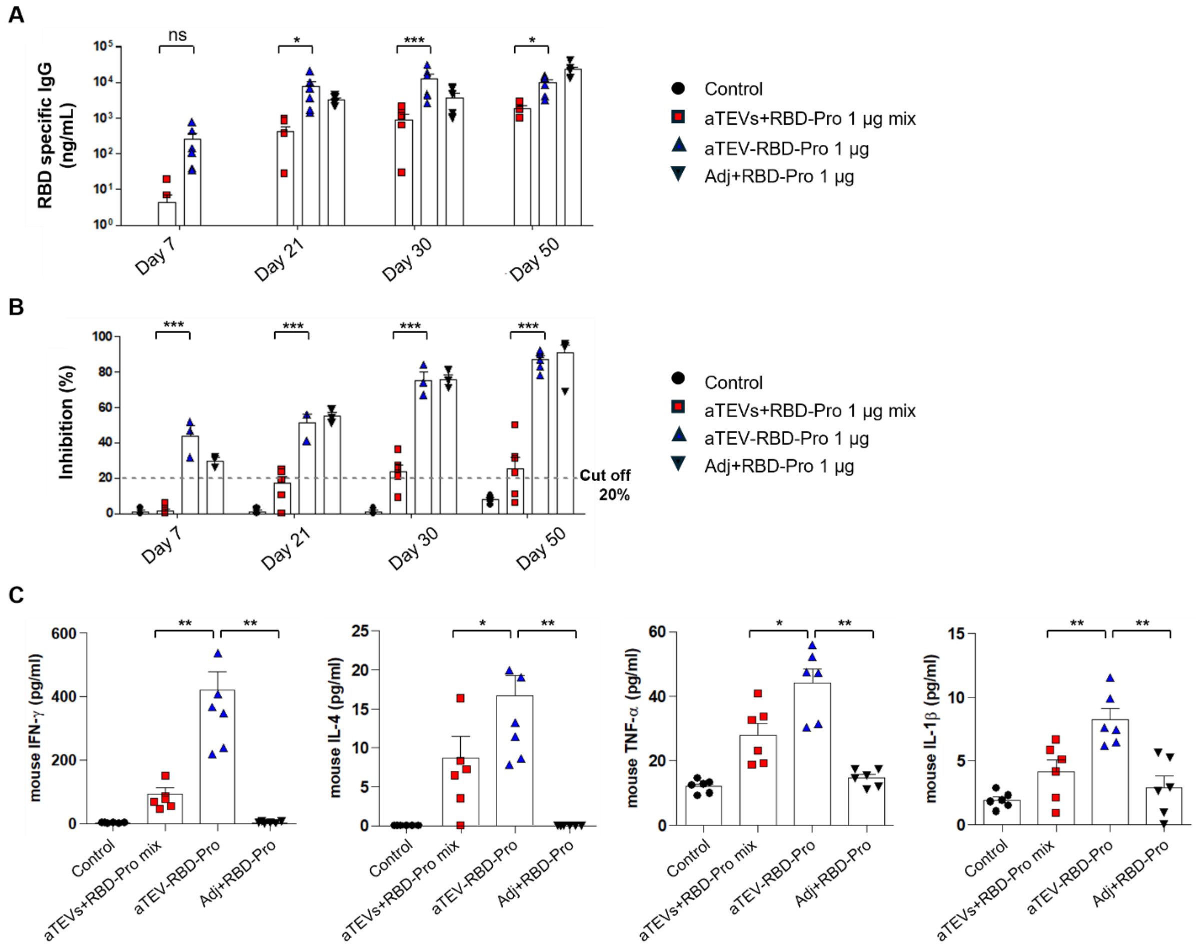


**Supplementary Figure 4.** Efficacies of antigen-encapsulated EVs and a simple mixture of antigen and EVs. BALB/c mice (n = 6/group) were immunized on days 0, 14, and 21 with a mixture of aTEVs and RBD (aTEVs+RBD mix), aTEV-RBD-Pro, or alum-adjuvanted RBD-Pro (Adj+RBD-Pro). (A) Level of RBD-specific IgG in plasma at days 7, 21, 30, and 50. (B) Level of neutralizing antibodies in plasma as determined by SARS-CoV-2 surrogate virus neutralization test at days 7, 21, 30, and 50. (C) Cellular immune response as determined in a cytokine assay in splenocytes. ns, non-significant; *P < 0.05, **P < 0.01, ***P < 0.001.

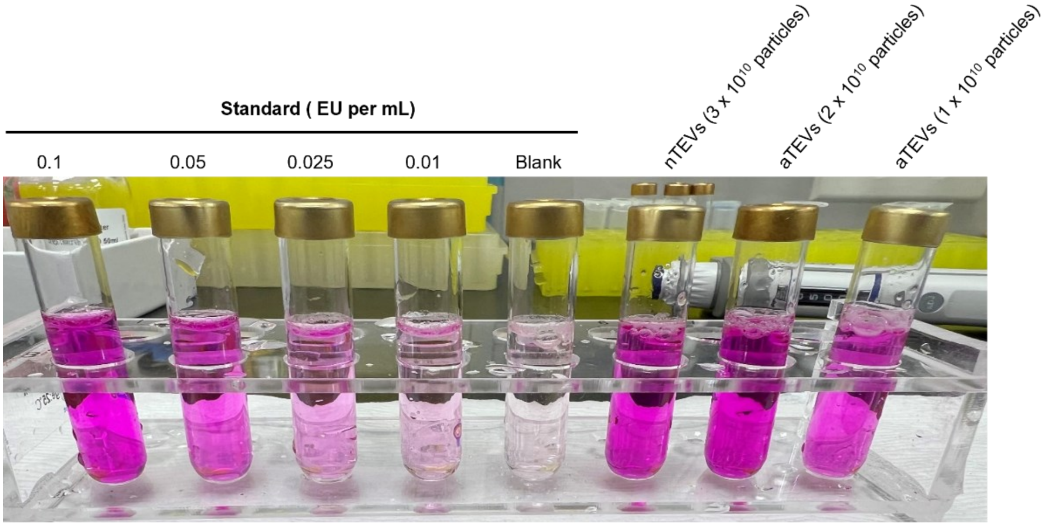


| | Endotoxin levels per $10^{10}$ particles |
|---|---|
| nTEVs | 0.03 $\pm$ 0.009 EU |
| aTEVs | 0.07 $\pm$ 0.003 EU |

**Supplementary Figure 5**. Detection of residual LPS levels in naïve TEVs (nTEVs) and activated (aTEVs). Residual LPS levels in nTEVs and aTEVs were measured using an endotoxin assay to confirm that the immunostimulatory effects of aTEVs were not due to LPS contamination.
